# The sphingosine-1-phosphate receptor functional antagonist fingolimod exacerbates neuroinflammation and tau pathology in a mouse model of tauopathy

**DOI:** 10.64898/2026.09.01.748583

**Authors:** Michelle D Rudman, Alexandra Litvinchuk, Cathryn Smith, Javier Remolina Serrano, Melissa Manis, Xin Bao, Carla M. Yuede, Jason D. Ulrich, David M Holtzman

**Author notes:** Correspondence: David M. Holtzman.

## Abstract

**Objective:** T cells are increased in the brain in Alzheimer’s disease (AD) and primary tauopathies where they correlate with tau pathology. In mouse models, T cells exacerbate tau-related neuroinflammation and neurodegeneration, and depletion of T cells is neuroprotective. Here, we investigate whether the multiple sclerosis drug fingolimod may inhibit infiltration of T cells into the brain in the setting of tau pathology and thereby attenuate tau-mediated neurodegeneration.

**Methods:** P301S-tau transgenic mice expressing human APOE4 (TE4), which develop tau pathology, neurodegeneration, and neuroinflammation with T cell accumulation, were randomly assigned to receive 1 mg/kg/d fingolimod in drinking water or standard mouse drinking water beginning just prior to the accumulation of neurofibrillary tangle pathology at 6 months of age. Mice were then assessed with a battery of cognitive behavioral assays at 9 months of age and euthanized at 9.5 months of age to analyze the blood and brain using flow cytometry, single molecule array (SIMOA) technology, stereotactic brain measurements, and immunohistochemistry.

**Results:** Fingolimod produced the expected lymphopenia in peripheral blood but unexpectedly exacerbated brain T cell accumulation, microgliosis, astrogliosis, and tau pathology, with no significant effect on neurodegeneration or behavioral impairment in TE4 mice.

**Interpretation:** These results indicate fundamental differences in the nature of the adaptive immune response to tau pathology versus other neuroinflammatory diseases such as multiple sclerosis and underscores the importance of understanding the impact of immune modulating therapies on all aspects of AD-related pathology in preclinical models prior to consideration and initiation of clinical trials.

## INTRODUCTION

Alzheimer’s disease (AD) is characterized pathologically by extracellular amyloid-β plaques and intracellular neurofibrillary tangles as well as a robust immune response. The innate immune response modulates both amyloid and tau pathology, with several immune-related AD risk loci such as *TREM2* highlighting the importance of microglial function.^1^ Emerging evidence also implicates the adaptive immune system in AD pathogenesis. T cells are increased in the brain in AD, in primary tauopathies, and in mouse models of AD-related pathology, and correlate spatiotemporally with tau pathology.^2–6^ These T cells consist predominantly of CD8^+^ effector memory T cells and show evidence of clonal expansion, suggesting an antigen-specific response to tau pathology.^2, 7, 8^ This is supported by the recent finding that type 1 conventional dendritic cells, the professional antigen presenting cells for CD8^+^ T cells, are required for the CD8^+^ T cell response to tau pathology and that antigens from the brain can be presented to T cells by antigen presenting cells in the deep cervical lymph nodes in mouse models.^9^ In the P301S-tau mouse model of tauopathy, depletion of circulating T cells is neuroprotective, suggesting a deleterious role for T cells in tau-related neuroinflammation and neurodegeneration.^2^ However, it is unknown how T cells accumulate in the brain and whether therapeutic inhibition of T cell infiltration may be neuroprotective in the setting of tau pathology.

Multiple therapies blocking T cell infiltration into the CNS have been developed for relapsing-remitting multiple sclerosis (RRMS). Fingolimod is a sphingosine-1-phosphate (S1P) receptor (S1PR) functional antagonist and was the first oral therapy approved for MS.^10^ S1P is a bioactive phospholipid produced by endothelial cells and red blood cells, creating high concentrations in the vasculature that drive chemotactic egress of lymphocytes from the lymph nodes into efferent lymphatic vessels and the circulation. Fingolimod is a pro-drug that, upon phosphorylation by sphingosine kinases, serves as a functional analog of S1P to bind and promote degradation of 4 of the 5 S1PRs thereby sequestering lymphocytes in lymph nodes. Fingolimod readily crosses the blood-brain barrier, accumulates in the CNS, and can bind S1PRs expressed on all CNS cells.^11^ In mouse models of amyloidosis, fingolimod and related S1PR functional antagonists reduce microgliosis, astrogliosis, and amyloid pathology and rescue cognitive deficits.^12–14^ However, evidence is limited and conflicting regarding the effects of fingolimod on tau pathology. In the mixed amyloid-tau 3xTg mouse model, fingolimod reduced phosphorylated tau (p-tau) pathology, microgliosis, and behavioral deficits.^14^ Conversely, fingolimod treatment in P301S-tau mice increased brain CD8^+^ T cell accumulation and hippocampal atrophy.^15^ It remains unclear how tau pathology and tau-related neuroinflammation may be affected by S1PR antagonism.

Here, we report that S1PR functional antagonism with fingolimod unexpectedly exacerbates tau pathology and neuroinflammation with increased micro- and astrogliosis as well as enhanced brain T cell accumulation in the P301S-Tau;ApoE4 (TE4) mouse model of tauopathy. These findings highlight unique features of the adaptive immune response to pure tau pathology and suggest that in the setting of tau pathology, the S1P-S1PR axis may play a protective role that has yet to be fully elucidated.

## METHODS

### Mice

Mice containing the human P301S-tau mutant transgene driven by a mouse prion protein (Prnp) promoter (PS19tg; Stock No. 008169, Jackson Laboratories) were backcrossed to C57BL/6 mice (Stock No. 027, Charles River) for >10 generations and crossed with human APOE4 knock-in mice (C57BL/6 background) in which endogenous mouse *Apoe* was replaced by human *APOE* flanked by LoxP sites^16^ to generate APOE4-homozygous P301S-tau-hemizygous transgenic mice on a C57BL/6 background (TE4 mice). Mice were bred, maintained, and used in experiments in a specific pathogen-free (SPF) animal facility in accordance with the guidelines of the Division of Comparative Medicine under protocols approved by the Institutional Animal Care and Use Committee at Washington University School of Medicine. All studies were conducted in accordance with the United States Public Health Service’s Policy on Humane Care and Use of Laboratory Animals.

### Fingolimod treatment

#### Dose-response Experiment

Male 7-9.5 month-old E4 mice were treated with 0, 0.1, 0.5, 1, or 5 mg/kg body weight fingolimod (Sigma-Aldrich, Y0002055) or water control by daily oral gavage and then euthanized on day 6 within 24 hours after the final dose. Blood was collected by cardiac blood draw to analyze leukocyte populations by flow cytometry.

#### Chronic treatment of TE4 mice

cages were randomized at 6 months of age to receive either 1 mg/kg/d fingolimod or standard drinking water (vehicle control) until 9.5 months of age. Fingolimod stock solutions (1.5 mg/mL) were made fresh weekly and stored at 4°C protected from light until use, and water bottles were replaced twice weekly. Daily water consumption was ∼5 mL/d/mouse, mouse weight ∼0.03 kg, and working concentration 0.006 mg/mL.

### Tissue Collection and Processing

Mice were euthanized at 285 days (±3d) of age by intraperitoneal injection of pentobarbital (200 mg/kg). Blood was collected via cardiac blood draw and mice were perfused with 3 U/ml heparin in Dulbecco’s PBS (DPBS, Gibco, Cat# 14040133) at 7mL/min for 3 minutes. The brain was extracted and bisected sagittally. The left hemisphere was fixed in 4% PFA overnight and stored in 30% sucrose at 4°C until it was sectioned coronally (30μm thick) on a Leica SM1020R microtome and stored in cryoprotectant solution (0.2 M DPBS, 15% sucrose, 33% ethylene glycol) at −20°C. The right hemisphere was dissected, snap frozen, and stored at −80°C.

### Plasma Neurofilament Light Chain Measurement

Plasma NfL concentration was measured with NF-Light Advantage PLUS Reagent Kit on a Simoa HD-X analyzer (Quanterix) per manufacturer’s instructions.

### Brain Stereological Analysis

Volumes of the hippocampus, piriform/entorhinal cortex (P/E Ctx), and left lateral ventricle were estimated from a sample of 30μm thick coronal sections of the left hemi-brain spaced every 180 µm spanning the full length of the hippocampus. Sections were stained with 0.1% cresyl violet with 0.05% glacial acetic acid for 2 minutes at 60°C, dehydrated and destained in ascending concentrations of ethanol, cleared with xylene, and then coverslipped with Cytoseal60 (Electron Microscopy Sciences, Cat# 18006). Slides were imaged with the NanoZoomer 2.0-HT whole slide imaging system (Hamamatsu Photonics) at 20x magnification, and areas of interest were traced using NDP.view2 software (Hamamatsu Photonics, U12388-01). Volumes were quantified using the following formula: volume (mm^3^) = (sum of areas, mm^2^) ∗ 0.180 mm. To quantify the thickness of the dentate gyrus granular cell layer, the thickness at the midpoint of the dorsal limb of the granule cell layer of the dentate gyrus was averaged across three tissue sections spaced 180µm apart within the range of approximate Bregma coordinates -2.3 to -2.7 mm.

### Immunohistochemistry

For glial immunohistochemistry, two brain sections per mouse were systematically selected, washed in TBS, blocked with 2% donkey serum in TBS with 0.25% Triton X-100, and incubated with primary antibodies (Iba1 (Novus Biologicals, NB100-1028; 1:500), CD68 (Biolegend, 137002; 1:250), Clec7a (Invivogen, mabg-mdect-2; 1:50), Tmem119 (Cell Signaling, 90840S; 1:500), GFAP (Invitrogen, 14-9892-82; 1:1000), MHC II (CST, 86285; 1:1000)) diluted in blocking buffer overnight at 4°C. Sections were then washed and incubated with fluorescent-conjugated secondary antibodies in blocking buffer for 1hr at room temperature (RT) the next day. Nuclei were stained with DAPI (Invitrogen, D1306; 1:1000 in TBS, 15 min), washed, and mounted in ProLong Diamond Antifade mounting medium (Invitrogen, P36961).

For AT8 staining, nonspecific peroxidase activity was blocked with 0.3% hydrogen peroxide in TBS for 10 min at RT followed by blocking of nonspecific antibody binding with 3% milk in TBS with 0.25% Triton X-100 (TBSX) for 30 min at RT. Sections were then incubated overnight at 4°C with 1:500 biotinylated AT8 antibody (Invitrogen, MN1020B) in blocking buffer, washed, and incubated in VECTASTAIN Elite ABC-HRP solution (Vector Laboratories, PK-6100) at RT for 1 hour. Finally, sections were stained in DAB solution (3,3’-diaminobenzidine (DAB) tablets, Sigma-Aldich, D5905-50TAB) with 30% hydrogen peroxide (Fisher Scientific,H325-100), washed, mounted on slides, and allowed to dry overnight. Sections were then dehydrated in increasing concentrations of ethanol followed by xylene and coverslipped with Cytoseal60 (ThermoFisher Scientific, 8310–16).

For T cell immunofluorescence, brain sections were washed with 0.1% TBSX and blocked with 10% normal donkey serum, 1% bovine serum albumin (BSA), and RecombiMAb anti-mouse CD16/CD32 FcR blocker (BioxCell, CP025; 1:400) in 0.5% TBSX for 2 hour at RT, followed by three days incubation at 4°C with primary antibodies [rat anti-CD3ε (Leinco Technologies, C2457 ; 1:400), human anti-CD4 (Miltenyi Biotec, 130-123-215; 1:50), rabbit anti-CD8α (Invitrogen, MA5-29682; 1: 200), and goat anti-IBA1 (Novus Biologicals, NB100-1028; 1:500) in 2% normal donkey serum and 0.2% BSA in 0.1% TBSX. Sections were then washed, incubated with fluorescent-labeled secondary antibodies (Invitrogen, 1:500) for 2 hours at RT, nuclei stained with DAPI, and sections mounted in ProLong Diamond antifade mounting medium.

### Image Acquisition and Analysis

For immunohistochemistry for glial markers and T cells, slides were imaged on a Leica Stellaris 8 Confocal Microscope with 20x objective and analyzed with ImageJ (NIH, v1.54k) and Imaris (Oxford Instruments, v10.2) software. For stereological analysis and quantification of tau pathology, slides were imaged with the Nanozoomer 2.0-HT system (Hamamatsu) and images were processed in ImageJ. Thresholding for percent area quantifications was set for each stain and background was subtracted by the ImageJ software before quantification. All analyses were performed blinded to treatment and genotype.

### Flow Cytometry

Blood aliquots were incubated in 5 mL ACK lysis buffer at RT for 5 minutes twice and filtered through a 70 µm cell strainer to remove RBCs. Cells were then incubated in LIVE/DEAD Fixable Blue Dead Cell Stain (Invitrogen, Cat no. L23105) per manufacturer’s protocol after which nonspecific antibody binding was blocked with 1:400 anti-mouse CD32/16 (Fc block, Leinco, Cat no. C247) for 10 minutes. Cells were then incubated in surface antibody cocktail with 1:400 Fc block in 2% FBS in DPBS for 30 minutes and washed in 2% FBS in DPBS. If no intracellular staining was utilized, samples were analyzed fresh. If intracellular staining was used, cells were fixed in 1x Fixation/Permeabilization Buffer (Foxp3/Transcription Factor Staining Buffer Set, eBioscience, Invitrogen, Cat No. 00-5523) for 40 minutes, washed twice in permeabilization buffer, and then incubated in intracellular antibody cocktail in permeabilization buffer for 40 minutes. A final wash was performed with permeabilization buffer and then cells were resuspended in PBS to run on a Cytek Aurora full spectrum flow cytometer. Unmixing and gating were performed in SpectroFlo software (Cytek). Antibodies are listed in Tables 1 and 2, with markers used to define cell populations described in Tables 3 and 4.

**Table 1.** Antibodies used in fingolimod dose-response blood flow cytometry.

|  | Marker | Clone | Fluorescent Tab | Supplier | Cat No. | Dilution |
| --- | --- | --- | --- | --- | --- | --- |
| 1 | CD45 | 30-F11 | Brilliant Violet 510 | BioLegend | 103138 | 1:200 |
| 2 | TCRβ | H57-597 | APC | BD Pharmingen | 553174 | 1:200 |
| 3 | CD4 | RM4-5 | Alexa Fluor 488 | BD Pharmingen | 557667 | 1:200 |
| 4 | CD8a | 53-6.7 | PE/Cyanine7 | BioLegend | 100722 | 1:200 |
| 5 | NK1.1 | PK136 | PE | BioLegend | 108708 | 1:200 |
| 6 | B220 | RA3-6B2 | PerCP-eFluor 710 | Invitrogen | 46-0452-82 | 1:200 |
| 7 | I-A/I-E | M5/114.15.2 | Brilliant Violet 421 | BioLegend | 101226 | 1:400 |

**Table 2.** Antibodies used for TE4 mice blood flow cytometry.

|  | Marker | Clone | Fluorescent Tag | Supplier | Cat No. | Dilution |
| --- | --- | --- | --- | --- | --- | --- |
| 1 | MHC II | M5/114.15.2 | APC | Cytek | 20-5321-U100 | 1:400 |
| 2 | CD8α | 53-6.7 | PE-Cy7 | Cytek | 60-0081-U100 | 1:200 |
| 3 | TCRβ | H57-597 | SuperBright600 | ThermoFisher | 63-5961-82 | 1:200 |
| 4 | CD45 | 30-F11 | BUV395 | ThermoFisher | 363-0451-82 | 1:200 |
| 5 | CD4 | REA604 | VioGreen | Miltenyi | 130-118-693 | 1:200 |
| 6 | CD19 | 1D3 | BUV615 | ThermoFisher | 366-0193-82 | 1:200 |
| 7 | TCR gd | GL3 | BV650 | ThermoFisher | 416-5711-82 | 1:200 |
| 8 | CD62L (L-selectin) | MEL-14 | PE | Cytek | 50-0621-U100 | 1:200 |
| 9 | CD44 | IM7 | SuperBright 780 | ThermoFisher | 78-0441-82 | 1:200 |
| 10 | CD3 | 17A2 | PacificBlue | BioLegend | 100214 | 1:200 |
| 11 | FOXP3 | FJK-16s | AlexaFluor 660 | ThermoFisher | 606-5773-82 | 1:200 |

**Table 3.** Defining markers for cell types for blood flow cytometry of fingolimod dose-response.

| Cell Type | Markers |
| --- | --- |
| Leukocytes | CD45<sup>+</sup> |
| T Cells | CD45<sup>+</sup>/ TCRβ<sup>+</sup>/NK1.1<sup>neg</sup> |
| CD8<sup>+</sup> T Cells | CD45<sup>+</sup>/ TCRβ<sup>+</sup>/NK1.1<sup>neg</sup>/CD8<sup>+</sup>/CD4<sup>neg</sup> |
| CD4<sup>+</sup> T Cells | CD45<sup>+</sup>/CD11b<sup>neg</sup>/TCRβ<sup>+</sup>/NK1.1<sup>neg</sup>/CD8<sup>neg</sup>/CD4<sup>+</sup> |
| NKT Cells | CD45<sup>+</sup>/CD11b<sup>neg</sup>/TCRβ<sup>+</sup>/NK1.1<sup>+</sup> |
| NK Cells | CD45<sup>+</sup>/CD11b<sup>neg</sup>/TCRβ<sup>neg</sup>/NK1.1<sup>+</sup> |
| B Cells | CD45<sup>+</sup>/CD11b<sup>neg</sup>/TCRβ<sup>neg</sup>/NK1.1<sup>neg</sup>/I-A/I-E<sup>+</sup>/B220<sup>+</sup> |

**Table 4.**
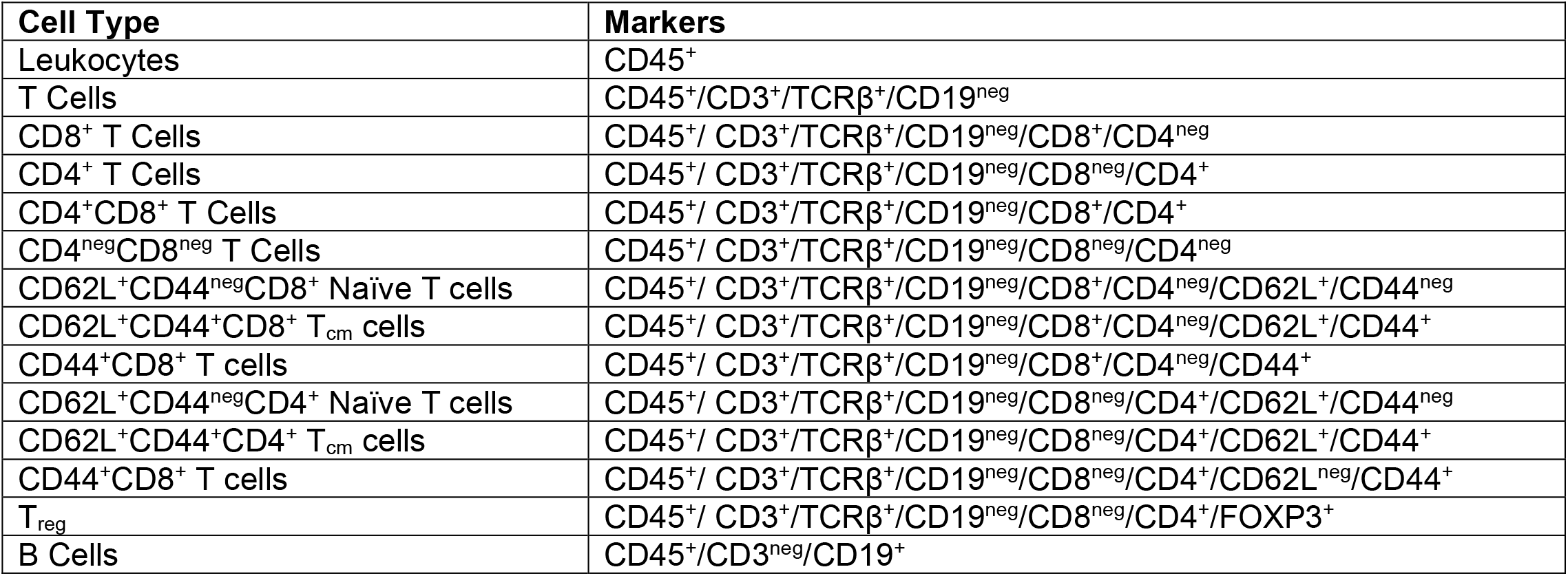
Defining markers for cell types for blood flow cytometry of TE4 mice.

### Nest Building Assay

Nest building assays were performed as described with minor modifications.^17^ 9.5 month-old mice were separated into clean cages with 3.0 g of nestlet material per cage and left overnight for about 20 hours. The morphology of the nest was scored and the total weight of unshredded nestlet pieces >0.1 g was recorded. Morphology scoring as follows: 1 = nestlet untouched, 2 = minimal shredding of nestlet, 3 = nestlet mostly shredded but dispersed across greater than one quadrant of the cage, 4 = nestlet mostly shredded and gathered within one quadrant of the cage, but with < 50% of the edges higher than the height of the mouse laying in the nest, 5 = most of nestlet material shredded, gathered into an identifiable nest, with >50% of the edges higher than the height of the mouse laying in the nest. Nestlet scoring was performed by a blinded scorer with confirmation by review of photos by a second scorer.

### Conditioned Fear Assays

Fear conditioning tests were performed in E4 and TE4 mice at 9 months of age as previously described.^18^ In brief, on day one each mouse was placed into the conditioning chamber and baseline freezing response was recorded for 2 min, after which a conditioned stimulus (80 dB tone) was presented for 20s followed by an unconditioned stimulus (1.0 mA electric shock) presented for 1s. This tone-shock (T-S) pairing was repeated during the last 20s of the next 2 min, and the freezing response quantified after each of the three tone-shock pairings. Twenty-four hours after training, each mouse was placed into the original chamber, and freezing behavior in response to contextual cues without tone or electric shock present was recorded for 8 min. Twenty-four hours after the contextual freezing response test, each mouse was placed into a new chamber to measure the conditioned response to the auditory cue. Baseline freezing behavior in the “altered context” was recorded for 2 min, followed by 8 min of tone presentation. FreezeFrame (Actimetrics) software was used for mouse behavior recording and analysis.

### Statistics

Most data were analyzed in GraphPad Prism 10 software and are presented as mean +/− SEM (* p<0.05, **p< 0.01, *** p <0.001, **** p<0.0001). For two-group comparisons, Student’s t-test was used. For multiple comparisons, ANOVA followed by appropriate post-hoc testing was utilized and is specified for each experiment in the figure legend. Nestlet scores were analyzed in the ordinal package in R 4.6.0 using a cumulative link model with genotype and treatment as fixed factors. Pairwise Tukey-corrected post hoc contrasts and corresponding p-values were calculated using estimated marginal means on the linear predictor scale in the emmeans package. All samples or animals were included in the statistical analysis, the n per group is included in the figure legends. Fear conditioning data was analyzed using Two-way Repeated Measures ANOVAs for differences between genotypes and treatment over time, and Two-way Factorial ANOVAs for total freezing behavior during each session. To control for multiple comparisons, we applied the False Discovery Rate (FDR) approach using the two-stage linear step-up procedure of Benjamini, Krieger, and Yekutieli, as implemented in GraphPad Prism, version 11.0.1. The desired FDR (Q) was set to 5% (0.05).

## RESULTS

### S1PR functional antagonism with fingolimod depletes circulating peripheral lymphocytes but increases T cells in the cortex and hippocampus of TE4 mice

To evaluate whether S1PR functional antagonism would reduce tau-related brain T cell accumulation, E4 and TE4 mice were treated with 1 mg/kg/d fingolimod in drinking water beginning at 6 months of age, an age where pathological tau phosphorylation is present without overt neurodegeneration or pronounced neuroinflammation or T cell accumulation.^2^ Fingolimod caused rapid and sustained lymphopenia at low doses with no appreciable off target effects on other leukocyte populations or toxicity with chronic treatment (Supplemental Figure 1, Figure 1A). In the blood, there was a shift in the composition of the small population of remaining lymphocytes consistent with fingolimod’s mechanism of action, with dramatic depletions of naïve CD4^+^ and CD8^+^ T cells and the majority of remaining T cells comprised of activated CD44^+^ T cells (Figure 1B). Significant decreases were also observed in circulating B cells, as anticipated, and regulatory T cells.

**Figure 1.**
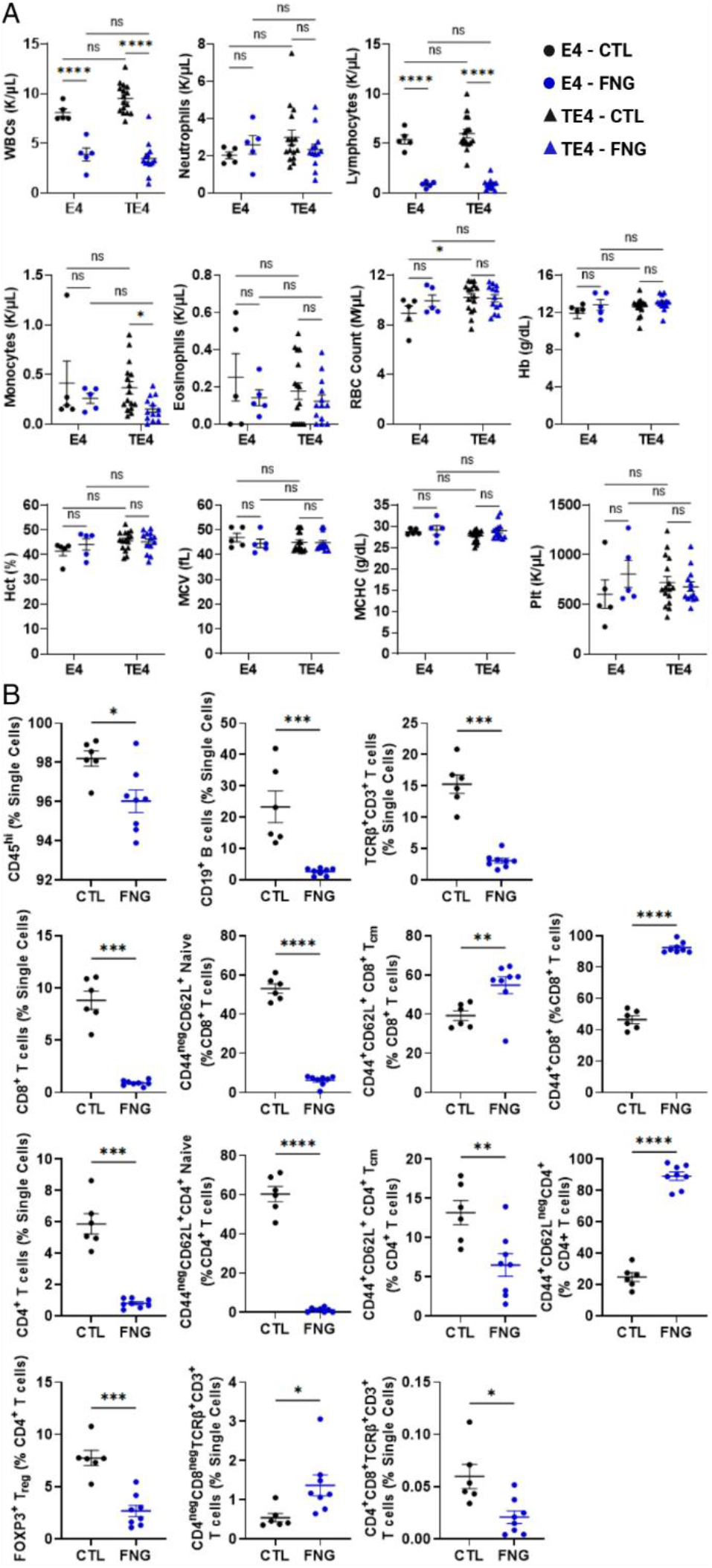
Fingolimod significantly reduces circulating lymphocytes in TE4 mice with preferential sequestration of B cells and naïve and central memory T cells and relative sparing of activated T cells. (A) Complete blood count from E4 and TE4 mice treated with water control or 1 mg/kg/d fingolimod in drinking water from 6 to 9.5 months of age. White blood cells (WBCs), red blood cells (RBCs), hemoglobin (Hb), hematocrit (Hct), mean corpuscular volume (MCV), mean corpuscular hemoglobin concentration (MCHC), and platelets (Plt). E4-CTL and E4-FNG N = 5, TE4-CTL N = 16, TE4-FNG N = 13. *p < 0.05, ****p<0.0001, Two-way ANOVA with Fisher’s LSD post-test, α = 0.05. (B) Flow cytometry of blood from 9.5-month-old TE4 mice treated with fingolimod from 6 to 9.5 months of age. Central memory T cells (T_cm_), regulatory T cells (T_reg_). E4 N = 6, TE4 N = 8. *p<0.05, **p<0.01, ***p<0.001, ****p<0.0001, Student’s t test, α = 0.05.

TE4 mice accumulate T cells in the brain in a manner that correlates spatiotemporally with the development of tau pathology and neurodegeneration, and the number of T cells in the brain is significantly increased compared to tau-negative age-matched control mice.^2^ We observed the expected increase in T cells in the cortex and hippocampus of control water-treated TE4 mice compared to tau-negative controls. Fingolimod treatment did not alter T cell numbers in the brain in tau-negative mice. In contrast to the significant depletion of circulating lymphocytes in the blood, however, TE4 mice treated with fingolimod had increased T cell accumulation in the cortex and hippocampus by 9.5 months of age compared to control water-treated mice (Figure 2). Increases were observed for both CD4^+^ and CD8^+^ T cell populations in the brain with no significant shift in the CD4^+^/CD8^+^ T cell ratio.

**Figure 2.**
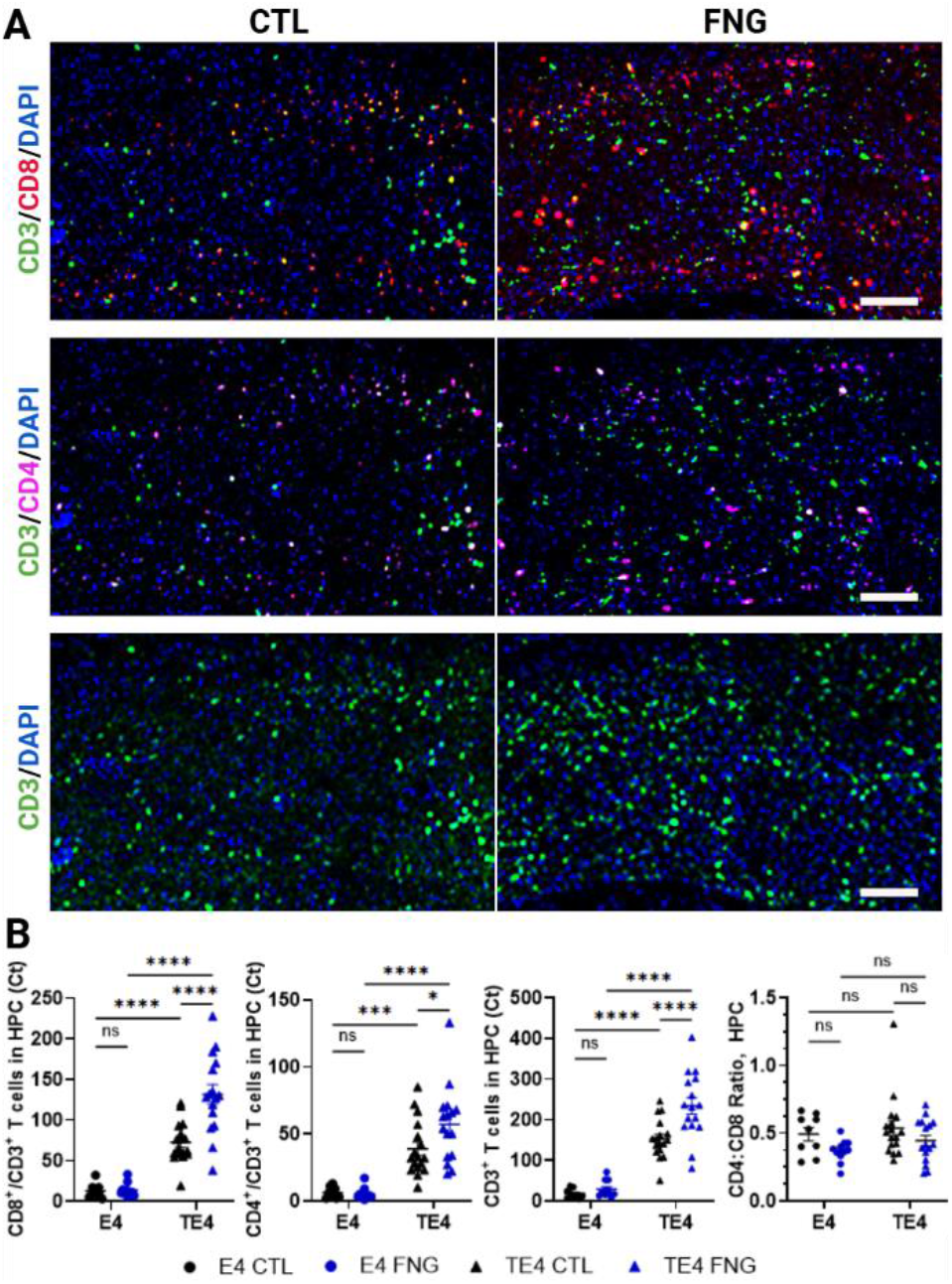
Chronic fingolimod treatment increases T cell accumulation in the hippocampus of aged TE4 mice. (A) Representative images of immunohistochemistry for T cell markers CD3, CD4, and CD8 in the hippocampus (HPC) of TE4 mice treated with control water (CTL) or fingolimod (FNG) from 6 to 9.5 months of age. Scale bar = 100 µm. (B) Quantification of number of T cells averaged from two sections of HPC per mouse from E4 and TE4 mice treated with CTL or FNG. E4-CTL N = 9, E4-FNG N = 11, TE4-CTL N=17, TE4-FNG N=16. *p<0.05, ***p<0.001, ****p<0.0001. Two-way ANOVA with Fisher’s LSD post-test, α = 0.05.

### S1PR functional antagonism with fingolimod exacerbates tau-related neuroinflammation in TE4 mice

In the setting of tau pathology, microglia and astrocytes adopt an activated phenotype characterized by the upregulation of genes associated with phagocytosis, lipid metabolism, and proinflammatory responses.^19^ Fingolimod treatment exacerbated microgliosis in the hippocampus and piriform/entorhinal cortex of 9.5-month-old TE4 mice, as evidenced by increased expression of Iba1 and activated microglial markers CD68 and MHC II (Figure 3A-C). There was also reduced expression of homeostatic marker Tmem119 and no significant change observed in the activated microglial marker Clec7a (Figure 3D, E). Similarly, reactive astrocytosis was increased by fingolimod treatment as evidenced by increased expression of GFAP in the piriform/entorhinal cortex compared to control-water treated TE4 mice (Figure 3F). These changes significantly correlated with the total number of T cells in the hippocampus (Figure 3G) and indicate increased neuroinflammation in the setting of increased T cells due to fingolimod.

**Figure 3.**
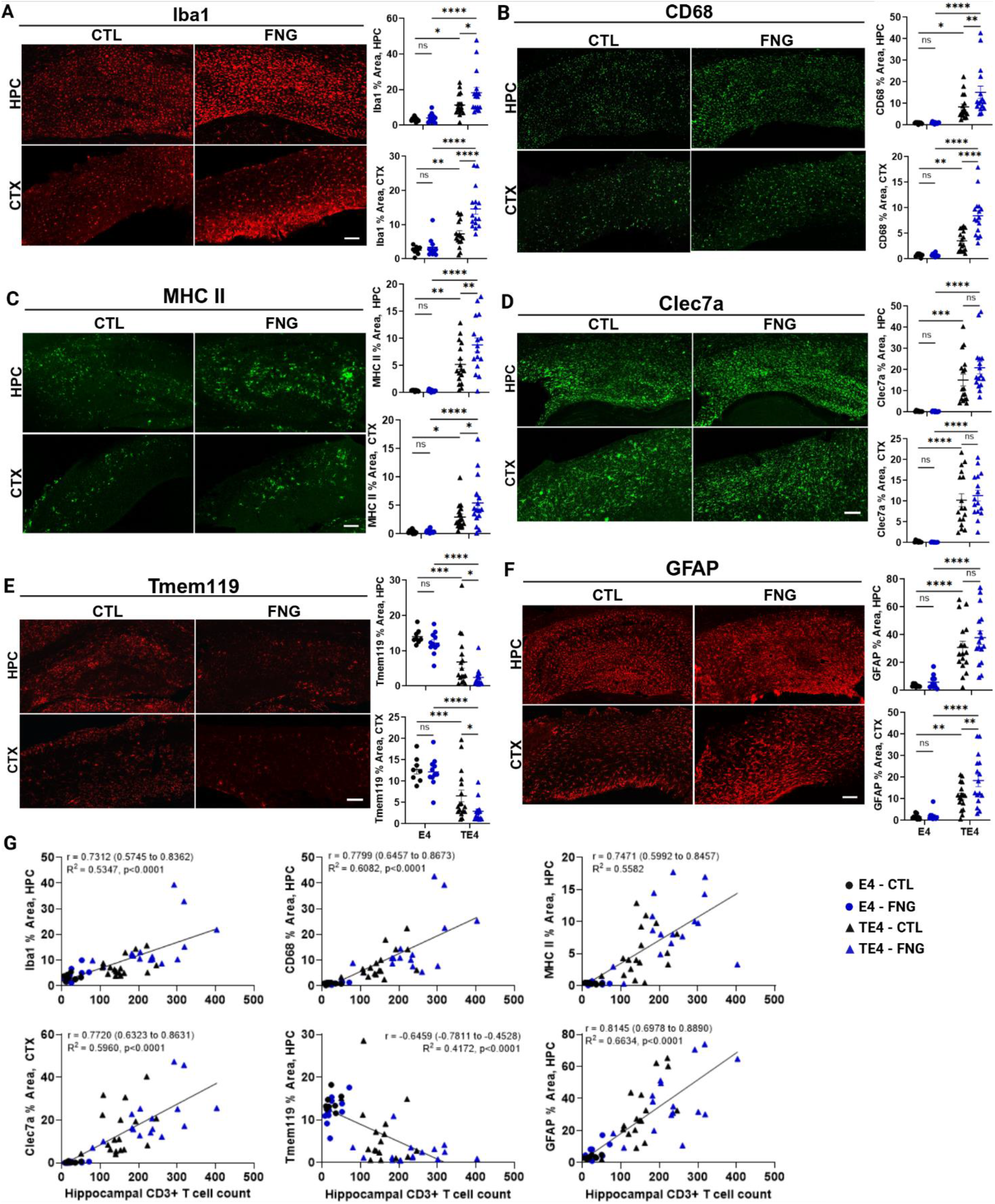
Fingolimod increases microgliosis and astrogliosis in aged TE4 mice. Representative images and quantification of percent area (% Area) positive from immunohistochemistry of hippocampus (HPC) and piriform/entorhinal cortex (CTX) for markers of micro- and astrogliosis including (A) Iba1, (B) CD68, (C) MHC II, (D) Clec7a, (E) Tmem119, and (F) GFAP. Scale bar = 100 µm. E4-CTL N = 9, E4-FNG N = 11, TE4-CTL N = 17, TE4-FNG N = 16. *p<0.05, **p<0.01, ***p<0.001, ****p<0.0001, Two-way ANOVA with Fisher’s LSD post-test, α = 0.05. (G) Correlation of hippocampal CD3+ T cell count with Iba1-, CD68-, MHCII-, Clec7a-, and Tmem118-positive percent area in hippocampus (HPC). Pearson correlation (r), (95% Cl), two-tailed, α = 0.05.

### S1PR functional antagonism with fingolimod exacerbates pathologic tau accumulation in TE4 mice but does not alter the degree of neurodegeneration or cognitive impairment

TE4 mice deposit pathologic misfolded and hyperphosphorylated tau protein in the hippocampus and piriform/entorhinal cortex around 5-6 month of age leading to neuronal dysfunction, neurodegeneration, and cognitive behavioral impairment by 9.5 months of age.^2, 20^ Fingolimod treatment beginning at 6 months of age exacerbated tau pathology in the hippocampus and piriform/entorhinal cortex of TE4 mice by 9.5 months of age as evidenced by increased AT8-positivity (Figure 4A, B). Further, the degree of tau pathology correlated significantly with the total number of T cells in the hippocampus (Figure 4C).

**Figure 4.**
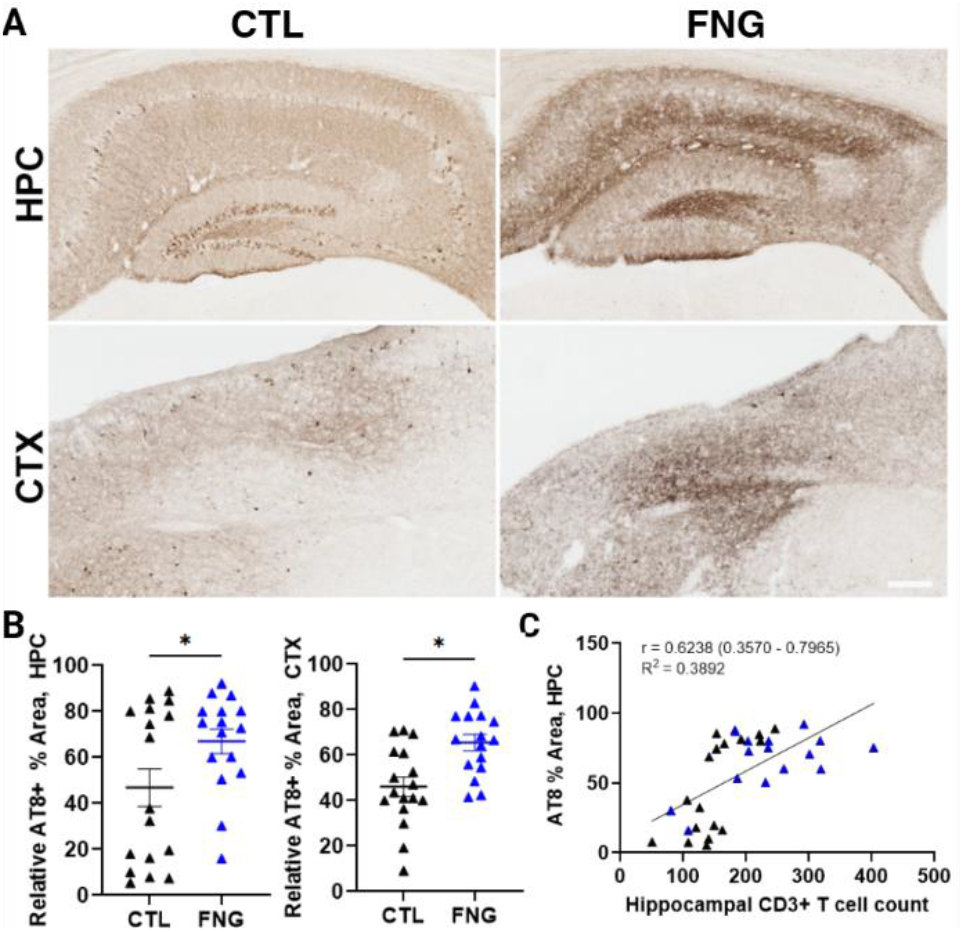
Fingolimod increases tau pathology in the cortex and hippocampus of aged TE4 mice. (A) Representative images of immunohistochemistry for phosphorylated tau (AT8) in hippocampus (HPC) and piriform/entorhinal cortex (CTX) of TE4 mice. Scale bar = 100 µm. (B) Quantification of percent (%) area AT8-positive. TE4-CTL N=17, TE4-FNG N=16. *p<0.05, Student’s t test, α = 0.05. (C) Correlation of hippocampal CD3+ T cell count with AT8-positive percent area in hippocampus (HPC) of TE4 mice. Pearson correlation (r), (95% Cl), two-tailed, α = 0.05.

Despite the increase in pathologic tau accumulation and neuroinflammation seen in fingolimod-treated TE4 mice, no significant differences were observed in measures of neurodegeneration or cognitive behavioral impairments. At 9.5 months of age, neurodegeneration was assessed by volumetric analysis of the hippocampus, piriform/entorhinal cortex, and ventricles. TE4 mice develop significant hippocampal and cortical atrophy by 9.5 months of age compared to tau-negative controls.^2^ Fingolimod treatment did not significantly alter the degree of brain atrophy in TE4 mice and fingolimod treatment also did not affect normal brain volumes in tau-negative controls (Figure 5A-B). Similarly, levels of plasma neurofilament light chain, a biomarker for neurodegeneration, were not significantly changed in fingolimod-treated TE4 mice (Figure 5C). Statistically significant correlations were observed between the number of hippocampal T cells and all measures of brain volumes suggesting that increased T cell accumulation may exacerbate tau-mediated neurodegeneration (Figure 5D).

**Figure 5.**
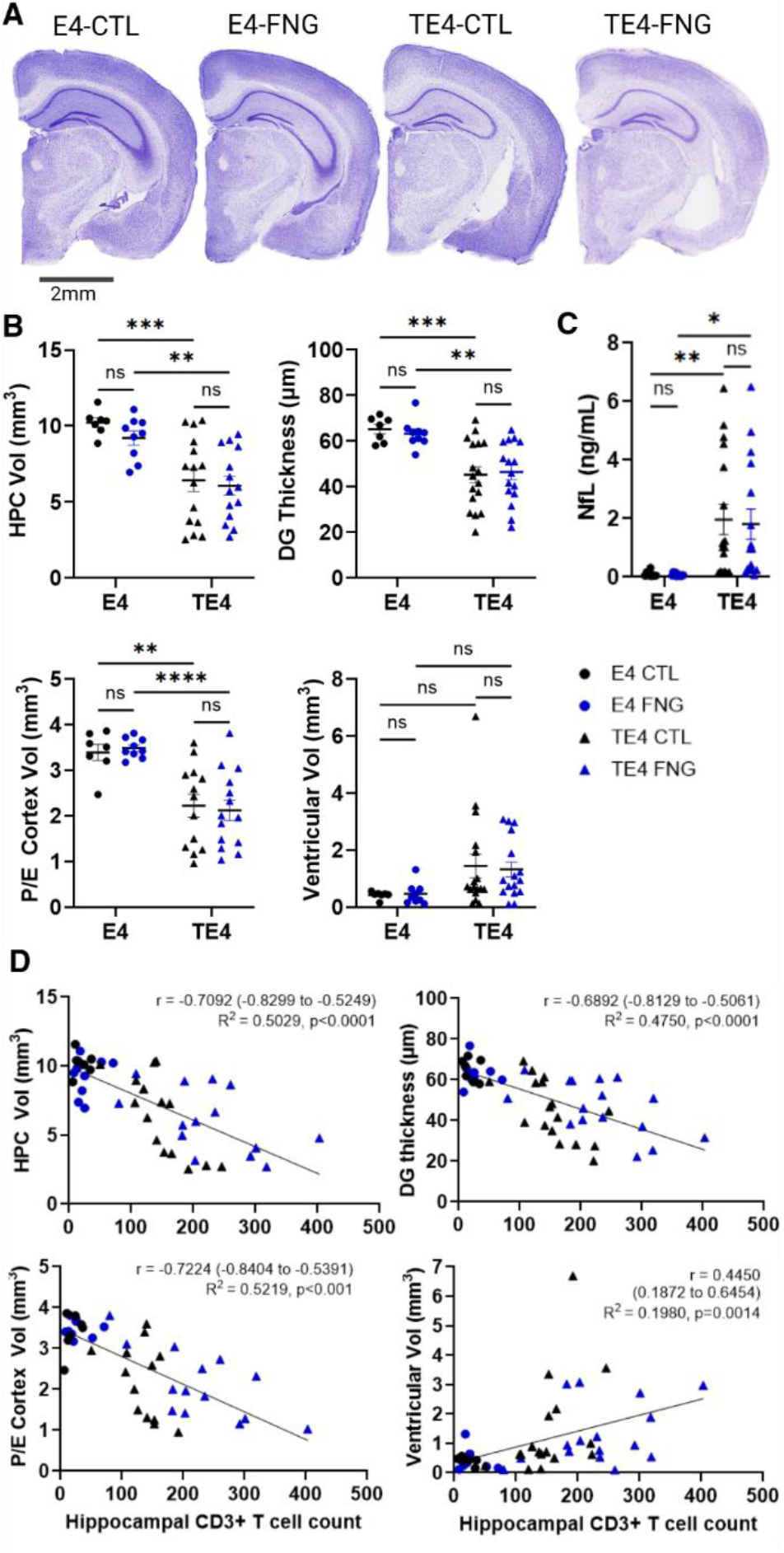
Fingolimod does not significantly alter neurodegeneration in aged TE4 mice though neurodegeneration correlates with hippocampal T cell accumulation. **(A)** Representative images of cresyl violet-stained left hemibrain. Scale bar 2mm. **(B)** Hippocampal volumes (HPC Vol, mm^3^), dentate gyrus (DG) thickness (µm), piriform/entorhinal cortex volumes (P/E Cortex Vol, mm^3^), and ventricular volumes (mm^3^) of E4 and TE4 mice treated with fingolimod (FNG) or control water (CTL) from 6 to 9.5 months of age. E4-CTL N = 7, E4-FNG N = 9, TE4-CTL N = 15-17, TE4-FNG N = 14-16. *p<0.05, **p<0.01, ***p<0.001, ****p<0.0001, 2-way ANOVA with Fisher’s LSD post-test, α = 0.05. **(C)** Plasma neurofilament light chain (NfL), pg/mL, E4 N = 9-12, TE4 N=16-17, *p<0.05, **p<0.01, 2-way ANOVA with Fisher’s LSD post-test, α = 0.05. **(D)** Correlation of hippocampal CD3^+^ T cell count with HPC volumes, DG thickness, P/E Cortex volumes, and Ventricular Volumes. E4-CTL N=7, E4-FNG N=9, TE4-CTL N=13-17, TE4-FNG N=14-16. Pearson correlation (r), (95% CI), two-tailed, α = 0.05.

Cognitive function in mice was assessed between 9 and 9.5 months of age with a battery of cognitive behavioral assays including a nest building assay and auditory and contextual fear conditioning. Nest building assays have previously been shown to be sensitive for detecting behavioral impairment in TE4 mice.^2^ TE4 mice showed the expected deficits in nest morphology and unshredded nestlet weight compared to tau-negative controls and fingolimod treatment did not alter nesting behaviors (Figure 6A). During tone-shock pairing for conditioned fear assays, there was a significant Time X Genotype effect [F(6,98) = 2.454, p = 0.0297] (Figure 6B). In the auditory cue conditioning assay, there were no significant differences in baseline freezing behavior but freezing during the presentation of the auditory cue showed a significant Time x Genotype interaction [F(21,343) = 1.669, p = 0.0338] with post hoc comparisons showing the E4-CTL group freezing more during minute 7 than all other groups (Figure 6C). In the contextual fear conditioning assay, there was a significant main effect of genotype [F(3,49) = 5.047, p = 0.004], indicating impaired hippocampal-amygdala function in TE4 mice compared to tau-negative controls. Post hoc comparisons indicated that both TE4 groups had impaired freezing compared to E4-CTL mice during minutes 2,3,4 & 5, and that the E4-FNG group was freezing more than the TE4-FNG group during minute 8 (Figure 6D). No significant differences between TE4-CTL and TE4-FNG mice were observed in either conditioned fear paradigm. However, statistically significant correlations were observed between the number of hippocampal T cells and both the nestlet score, unshredded nestlet weight, and contextual conditioning total freezing time, suggesting that increased T cell accumulation may exacerbate cognitive behavioral impairment due to tauopathy (Figure 6E).

**Figure 6.**
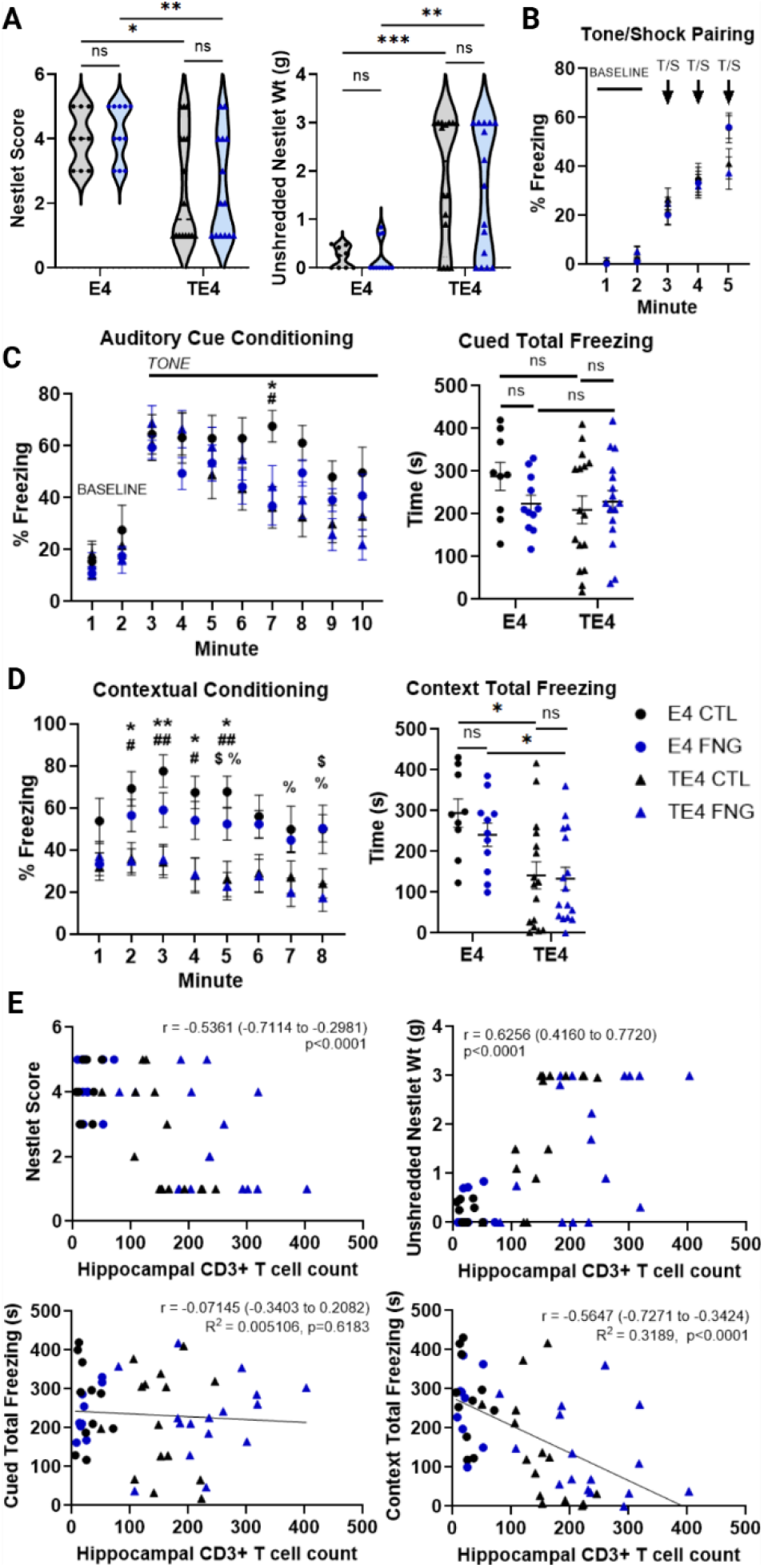
Fingolimod does not significantly alter behavior in aged TE4 mice though behavioral impairment correlates with hippocampal T cell accumulation. **(A)** Nestlet score and weight of the unshredded portion of a 3g nestlet (g), E4-CTL N = 9, E4-FNG N = 11, TE4-CTL N = 16, TE4-FNG N = 14, ordinal regression with Tukey post-test, α = 0.05. **(B)** Percent freezing per minute during tone/shock pairing and **(C)** auditory cue and contextual fear conditioning percent freezing per minute and total freezing time (s). E4-CTL N=9, E4-FNG N = 11, TE4-CTL N = 16, TE4-FNG N = 16. Two-way Repeated Measures ANOVAs for differences between genotypes and treatment over time, and Two-way Factorial ANOVAs for total freezing behavior during each session with the two-stage linear step-up procedure of Benjamini, Krieger, and Yekutieli (Q = 0.05). E4-CTL vs. TE4-CTL: *q < 0.05, **q < 0.01. E4-CTL vs. TE4-FNG: ^#^q < 0.05, ^##^q < 0.01. E4-FNG vs. TE4-FNG: *^%^*q < 0.05. E4-FNG vs. TE4-CTL: ^$^q<0.05. **(D)** Spearman correlation of hippocampal T cell counts with Nestlet score and Unshredded Nestlet Weight, and Pearson correlation(r) of hippocampal T cell counts with total freezing time in cued and contextual fear conditioning paradigms, E4-CTL N=9, E4-FNG N=11, TE4-CTL N=16, TE4-FNG N=16, (95% CI), two-tailed, α = 0.05.

## DISCUSSION

Increasing appreciation for T cell involvement in AD and primary tauopathies has sparked interest in the targetability of these pathways to ameliorate neurodegeneration and cognitive decline in AD. The S1P-S1PR pathway is known to play a critical role in trafficking lymphocytes to the brain in RRMS, where S1PR functional antagonism with fingolimod is effective for reducing recurrence of relapses and accumulation of disability.^10^ Recent studies also implicate S1P-S1PR signaling in promoting detrimental neuroinflammation, pathological amyloid accumulation, and cognitive impairment in the context of amyloid pathology.^12–14^ However, the role of S1P-S1PR signaling in the setting of tau pathology, which correlates more strongly with T cell accumulation, was unclear. One prior study reported that, in a model of pure tauopathy, short-term fingolimod treatment after the onset of tau pathology increases brain CD8^+^ T cell accumulation in a manner correlating with pS396- and AT8-positive p-tau pathology and exacerbates hippocampal neurodegeneration.^15^ This study was limited by small cohort size, shorter term treatment, and lack of analyses for glial phenotypes or behavioral correlates of hippocampal neurodegeneration. A second study using fingolimod in the mixed amyloid-tau 3xTg model demonstrated improvements in cognitive performance, reduced hippocampal Iba1^+^ cell density, and reduced AT8-positive tau pathology, but this was not clearly tied to effects on T cells in the brain as T cells were not increased in the brain in this model.^14^ Further it is not clear whether the observed reduction in tau pathology and microgliosis by fingolimod was related directly to tau pathology or indirect due to effects on amyloid pathology.

Our study builds on prior literature by demonstrating that fingolimod exacerbates tau-driven CD8^+^ and CD4^+^ T cell accumulation in the brain and increases micro- and astrogliosis as well as increased AT8^+^ tau accumulation in the hippocampus and piriform/entorhinal cortex. These results indicate that S1PR functional antagonism with fingolimod exacerbates tau pathology and tau-related neuroinflammation and T cell accumulation. While we did not observe exacerbated tau-mediated neurodegeneration or behavioral impairments with fingolimod, T cell numbers correlated with both neurodegeneration and behavioral impairment suggestive of a link between T cells and tau-dependent damage. Our ability to detect changes in neurodegeneration and behavioral impairments may be limited by the high variability of neurodegeneration observed in this TE4 cohort. This discrepancy may also indicate that fingolimod has direct protective effects on neurons, mitigating its deleterious effects on tau pathology and neuroinflammation. Prior literature has suggested, for example, that fingolimod may reduce NMDA receptor-mediated excitotoxic neuronal death.^21^ Future investigation into cell-specific effects of fingolimod in the context of tau pathology may further clarify this possibility.

The therapeutic effects of fingolimod in RRMS are primarily attributed to sequestration of activated lymphocytes in lymph nodes. However, the effects of fingolimod on lymphocyte subpopulations vary depending on the responsiveness of the subpopulation to S1P-S1PR signaling. In RRMS, there is robust infiltration of circulating T cells into active lesions with blood-brain barrier (BBB) breakdown. Central memory and naïve T cells are preferentially sequestered by treatment with fingolimod and fingolimod reduces the incidence of relapses, therefore implicating these populations in relapse pathogenesis. In contrast, secondary- and primary-progressive multiple sclerosis (SPMS/PPMS) are characterized by chronic accumulation of lymphocytes without significant BBB breakdown, and fingolimod is ineffective for reducing disability progression in PPMS^22^, suggesting that this chronic smoldering inflammation is fundamentally different. One key difference may lie in the subpopulations of lymphocytes that mediate these different responses. In SPMS/PPMS, parenchymal T cells acquire a tissue-resident memory phenotype, losing S1PR1 and CCR7 expression.^23^ Similarly, T cells from the cerebrospinal fluid of individuals with AD and the brains of mice with AD-related pathology consist primarily of CD8^+^ effector memory T cells.^2, 7^ CD8^+^ effector memory T cells are spared from fingolimod-mediated lymph node sequestration and become the dominant circulating T cell population in individuals treated with fingolimod due to the lack of CCR7-mediated retention in the lymph nodes. Likewise, tissue-resident memory T cells downregulate S1PRs as part of the transcriptional program that establishes tissue residency, and would not be expected to be significantly affected by S1PR-modulating treatments.^24^ TCR restimulation in tissues can also suppress S1P-mediated trafficking and could contribute to T cell accumulation in the brain, independent of fingolimod S1PR functional antagonism.^25^ The chronic inflammation associated with tau-related neurodegeneration may therefore be resistant to S1PR functional antagonism similarly to that of progressive forms of MS due to involvement of T cell subtypes that downregulate S1PRs.

While this may explain the inability of S1PR functional antagonism to suppress tau-related T cell accumulation in the brain, it remains unclear why T cell accumulation is increased. Potential mechanisms may include inhibition of T cell egress from the brain, inhibition of local T cell apoptosis, relative inhibition of protective/anti-inflammatory lymphocyte subpopulations, preferential sparing of more cytotoxic T lymphocytes, and/or direct effects in the CNS that exacerbate tau pathology and/or neuroinflammation. Fingolimod accumulates preferentially in the CNS in a time- and dose-dependent manner and could contribute to retention of lymphocytes in the brain, though this has not been demonstrated and would be challenging to distinguish from inhibition of apoptosis^26^. While total circulating lymphocyte populations are significantly decreased by fingolimod, the remaining circulating lymphocytes have been shown to retain or even have enhanced effector function, and the effect on regulatory T cells in the setting of tauopathy remains unknown.^27^

Regarding direct effects of fingolimod in the CNS, fingolimod and other S1PR functional antagonists attenuate LPS-induced chemokine release in astrocytes and microglia *in vitro* and suppress glial activation and expression of proinflammatory cytokines and chemokines when given concurrently with cuprizone in a mouse model of autoimmune-independent demyelination.^28–31^ In the experimental autoimmune encephalitis mouse model of MS, fingolimod not only attenuates disease activity, but astrocytic expression of S1PR1 is specifically required for the beneficial effects of fingolimod in this model indicating that astrocytes play a critical role in the therapeutic effects of S1PR antagonism in the setting of acute CNS autoimmune disease.^32^ Even in mouse models of chronic progressive CNS neuroinflammatory disease meant to mimic aspects of SPMS, fingolimod suppressed expression of proinflammatory cytokines and chemokines by astrocytes and microglia and attenuated clinical disease progression.^31^ By contrast, fingolimod does not ameliorate cuprizone-induced oligodendrocyte death and may actually exacerbate astrogliosis when the start of treatment is delayed.^30, 33^ This indicates opposing context- and timing-specific effects of S1PR antagonism, at least in the setting of toxin-induced demyelination, though detrimental effects have not been observed with delayed initiation of fingolimod in other disease models such as EAE.^34^ Potential etiologies for the increase in micro- and astrogliosis observed with fingolimod treatment in the setting of tauopathy may include secondary effects due to increased T cell accumulation. This is supported by the observed increase in MHC II expression with fingolimod treatment. MHC II is upregulated by IFNγ which is primarily produced by T cells, suggesting a link between T cell accumulation and the increase in gliosis in the setting of fingolimod treatment. Other potential contributors to increased gliosis include fingolimod-mediated exacerbation of tau pathology as well as differences in S1PR subtype expression on glial cells in the setting of tauopathy, resulting in different downstream effects compared to other chronic inflammatory diseases. Future studies examining tau-related changes in cell-specific S1PR expression profiles as well as direct effects of fingolimod on neuronal physiology and pathologic tau accumulation may help clarify whether these mechanisms contribute to the exacerbation of tau pathology and neuroinflammation by fingolimod.

Recently there have been growing calls for clinical trials to assess the efficacy of S1PR functional antagonists in AD based largely on preclinical data from mouse models of amyloidosis.^35^ AD, however, consists of multiple and complex pathologies and it is important to assess the potential effects of any new therapeutic on each of these various pathologies. This study provides evidence that S1PR functional antagonism exacerbates tau pathology and tau-related neuroinflammation in the setting of pure tauopathy. Particularly in the setting of symptomatic AD where tau pathology and neurodegeneration are already well-established, S1PR functional antagonists may therefore have the potential to exacerbate disease. It remains unclear what effects S1PR functional antagonism may have on tau pathology in the setting of mixed amyloid-tau pathology. Additional studies utilizing models of combined amyloid-tau pathology that develop neurodegeneration would therefore be of the utmost importance prior to clinical trials.

Clinical data is lacking on the risk of AD and primary tauopathies with chronic treatment with S1PR functional antagonists. Studies evaluating risk of AD in individuals with MS and those on immunomodulatory therapies have historically been limited by confounds from dementia due to MS and other co-pathologies in living participants making postmortem pathological diagnoses essential in these cases. However, with the recent advent of sensitive and specific biomarkers for AD, this question may now be more accurately assessed in living individuals as evidenced by emerging studies indicating that AD biomarker positivity is actually reduced in individuals with MS.^36, 37^ The use of plasma biomarkers in particular has enabled larger studies which will continue to provide new insights.^36^ Application of widely-available AD biomarkers in studies addressing risk of amyloid and tau pathology in those on immunomodulatory therapies could provide significant insights into these questions.

## Acknowledgements

This study was supported by grants from NIH AG085374 (DMH, JU), Rainwater Foundation (DMH), GHR Foundation (DMH), a gift from Ronald Shaich (DMH), grants from the Alzheimer’s Association (MDR), NIH R25 (MDR), and Knight ADRC REC Scholars program (MDR).

## Author Contributions

MDR, JDU, and DMH contributed to the conception and design of the study; MDR, AL, CS, JRS, MM, XB, CMY, JDU, and DMH contributed to the acquisition and analysis of data; MDR, AL, CMU, JDU, and DMH contributed to drafting the text or preparing the figures.

## Data Availability

The data that support the findings of this study are openly available in the WashU Medicine’s Digital Commons Data@Becker Repository with the following DOI: https://doi.org/10.17632/f3zf9rnc7n.

