## Supplemental Figure 1 for "The sphingosine-1-phosphate receptor functional antagonist fingolimod exacerbates neuroinflammation and tau pathology in a mouse model of tauopathy"

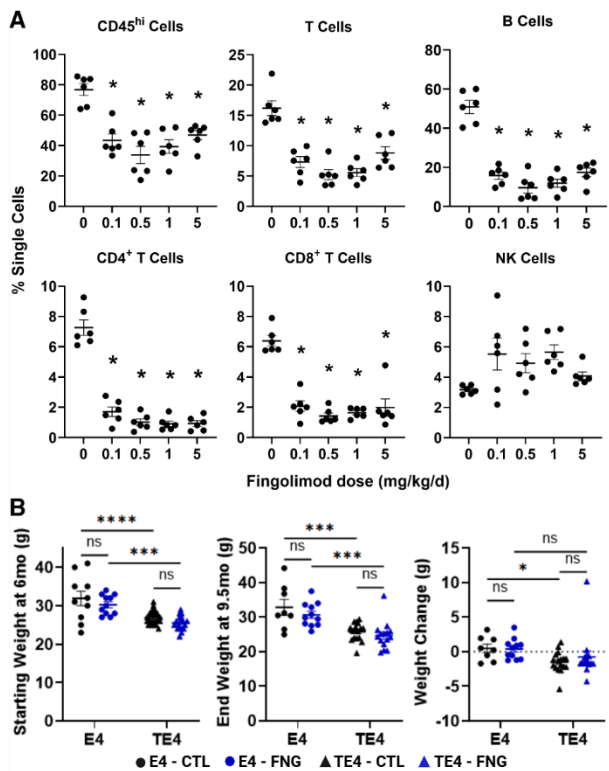

**Supplemental Figure 1. Oral fingolimod results in rapid depletion of circulating lymphocytes at low doses with no observed toxicity.** (A) Flow cytometry for leukocytes in peripheral blood samples of 7 - 9.5-month-old male E4 mice treated with one of 5 doses of fingolimod by daily oral gavage for 5 days. N = 6. \*p < 0.05 compared to vehicle control, remaining comparison ns, One-way ANOVA with Tukey post-test, $\alpha = 0.05$. (B) Starting and ending body weight and weight change (g) of E4 and TE4 mice treated with fingolimod in drinking water from 6 to 9.5 months of age. \*p<0.05, \*\*\*p<0.001, \*\*\*\*p<0.0001, Two-way ANOVA with Fisher's LSD post-test, $\alpha = 0.05$. E4-CTL N = 8-10, E4-FNG N = 12, TE4-CTL N = 17-21, TE4-FNG N = 15-18.
